# A Dual-Gate OECT Platform for Drift-Suppressed and High-Sensitivity Real-Time Biosensing

**DOI:** 10.64898/2026.09.10.750357

**Authors:** Stefany J. Kissovsky, Scott Keene, George G. Malliaras, Gabriele S. Kaminski Schierle

## Abstract

Organic electrochemical transistors (OECTs) enable high transconductance at low operating voltages through the volumetric charging of organic mixed ionic/electronic conductors (OMIECs), making them attractive transducers for biosensing. However, the same ionic -electronic coupling makes OMIECs, such as poly(3,4-ethylenedioxythiophene)-poly(styrenesulfonate) (PEDOT:PSS), sensitive to different ions and various environmental fluctuations, so drift and memory effects may compromise the quantitative readout in complex biofluids. In this work, we introduce a novel OECT-based sensor configuration that minimises drift by addressing a single channel with two gates: a functionalised sensing gate and a reference gate that allows real-time decoupling of environmental and channel-state fluctuations. A pulsed gate-biasing protocol is introduced to promote channel conductivity recovery and mitigate state accumulation. Using enzymatic glucose sensing as a proof of concept, the platform suppresses time-dependent drift by ∼96%, eliminates temperature-induced variations up to 99% across the 20-70 ^₀^C range, improves calibrated sensitivity by 55% and reduces interferent-induced deviations under physiologically relevant conditions. By improving both signal stability and analytical reliability, this strategy addresses a central limitation of OECT biosensors and advances their use in high-sensitivity point-of-care biosensing in complex biological environments.

## 1. Introduction

Point-of-care testing (POCT) has grown significantly over the past decade, driven by the increasing demand for rapid and reliable health monitoring through the analysis of patient samples containing biomarkers within complex physiological matrices.^1–3^ Organic electrochemical transistors (OECTs) have emerged as promising platforms for POCT for biosensing due to their biocompatibility, high amplification and high sensitivity as electrolyte-gated transistors. OECT-based platforms have also been reported to outperform conventional electrochemical sensing in metrics such as limit of detection, stabilization time, and signal-to-noise ratio.^8^ Unlike organic field-effect transistors (OFETs) and electrolyte-gated organic field-effect transistors (EGOFETs), where gating modulates only the charge at the channel-electrolyte surface, OECTs enable bulk ion-electron coupling and consequently much higher transconductance at lower bias, typically below 1 V. ^4-7^ This is mainly due to the use of organic mixed ionic-electronic conductors (OMIECs), commonly used for channel materials in OECTs. They enable volumetric ion-electron coupling, delivering high sensitivity while supporting operation in high-ionic-strength aqueous electrolytes relevant to biofluids.^4,5,9^

OMIECs are also history-dependent materials whose conductivity depends on ionic content and redox state. In enzymatic biosensor OECTs, the measured output is the drain current of the OMIEC channel, implicitly assuming that changes in channel conductance primarily track the analyte-driven electrochemical process.¹⁰ In practice, drift can arise from coupled redox and environmental effects, i.e. slow OMIEC redox kinetics together with changes in ambient oxygen and humidity can modulate the channel’s redox/ionic state and thus its effective doping level.^11^ Moreover, in a polarized channel, ionic swelling could modulate the ions penetrating the channel and therefore lead to an apparent degradation of channel conductivity.^12,13^

Poly(3,4-ethylenedioxythiophene)-poly(styrenesulfonate) (PEDOT:PSS) has been one of the most widely used channel materials due to its good solubility and high ion-permeability that facilitates mixed ionic–electronic transport. However, its conductance is highly sensitive to fluctuations in pH, humidity, and temperature.^14–16^ OECTs with PEDOT:PSS and composite channels can exhibit intrinsic memory characteristics due to ion entrapment, an effect intentionally exploited for electrochemical neuromorphic organic devices (ENODs).^17–19^ Thus, PEDOT:PSS channel-state drift and history dependence can masquerade as analyte response in complex biofluids, compromising a reliable enzymatic readout.

To improve selectivity and suppress interferent-driven artefacts, state-of-the-art enzymatic OECT biosensing has largely focused on designing the gate/electrolyte interface to localize the enzymatic reaction at the electrode. As a result, limits of detection as low as 10 nM have been reported with selective membranes such as Nafion–graphene oxide composites, and gate-modification strategies, such as molecularly imprinted polymer-modified gates and fiber-integrated sensors, for analytes including lactate, ascorbic acid, and glucose.^7,20,21^ However, these advances predominantly optimize the gate interface specificity, while the OMIEC channel remains susceptible to environmental and operational variations, motivating referencing and multi-electrode architectures that can explicitly correct for channel-state changes during sensing.

To minimize drift-induced artefacts, recent OECT biosensing strategies have explored paired-device and multi-gate architectures intended to provide a more reliable readout. Differential Wheatstone-bridge configurations using matched OECT pairs can suppress interference from electro-oxidizable species with good sensitivity, but impedance mismatch and device-to-device variability between the two channels can introduce noise.^22^ A dual-gate configuration has been shown to successfully extend the biochemical reaction driving redox potential, protecting the channel from high potentials, yet without correcting for channel-state evolution during operations.^23^ A dedicated reference channel, functionalized for nonspecific interactions, has been incorporated by Harris et al. for subtraction of interferent adsorption and physical changes at sensing interfaces.^24^ This strategy effectively suppresses interface-driven baseline shifts; however, as with subtraction-based referencing, performance depends on the extent to which drift remains sufficiently a common-mode across the two channels despite intentionally different surface chemistries and associated interfacial kinetics. Song et al. further demonstrates that even under constant conditions, time-dependent drain-current drift evolves due to ion adsorption within the gate material and sensing layer in physiologically relevant matrices.^25^ Their dual-gate approach mitigates the observed drift and increases sensitivity but does not explicitly address drift originating from the OMIEC channel ion uptake and memory. Collectively, these approaches improve selectivity, drift tolerance, or operating range, yet they typically do not address dynamic channel-state evolution during enzymatic sensing, including ionic memory, doping dynamics, and time-dependent faradaic byproducts. These limitations motivate a self-referencing architecture that combines real-time monitoring of channel-state evolution with in situ channel reconditioning.

In this work, we introduce a shared-channel, self-referencing dual-gate OECT platform that combines shared-channel drift monitoring with pulsed channel reconditioning to separate analyte-dependent signals from channel-state fluctuations. The in-plane reference provides real-time channel-state calibration while a functionalized sensing gate performs biochemical transduction. The device is operated using a sense–replenish–ground control cycle: during the sense phase, analyte-driven modulation is recorded; during the replenish phase, the channel is actively reconditioned to promote recovery of channel conductivity and to reduce ion accumulation; and during the ground/reset phase, the gates are returned to 0 V to reduce residual surface charge and bring the device closer to its initial baseline. The internal reference provides a measure of common-mode drift and baseline shifts, while a model-based calibration factor compensates for device-to-device variability and interfacial kinetic mismatches, enabling analyte-dependent responses to be separated from environmental drift. Conceptually, the approach performs operando material-state calibration of PEDOT:PSS, turning its state evolution into a controlled variable.

Using enzymatic glucose sensing as a proof of concept, we evaluate this strategy under controlled environmental perturbations, and using physiologically relevant interferants, and complex biological media. Compared with standard continuous enzymatic OECTs, the proposed dual-gate platform stabilizes the OMIEC channel state by decreasing drift with ∼96% after 40 min and minimises temperature perturbations to ∼1% in the range of 20 -70 ^₀^C. The sense-replenish-ground gate bias operation tracks in real-time non-linear channel-state drifts during increasing glucose concentrations and actively enhances recovery of the PEDOT:PSS channel. Under glucose sensing with common interferents, the dual-gate platform further reduces interferent-induced drift by ∼72–97% relative to the raw response with the strongest suppression observed for ascorbic acid. By linking enhanced sensitivity with improved analytical reliability, this platform addresses a key barrier to OECT-based point-of-care diagnostics and advances real-time biochemical monitoring for healthcare applications.

## 2. Results and Discussion

### 2.1. Device structure

A shared-channel dual-gate organic electrochemical transistor (OECT) system, containing one sensing and one reference gate electrode, was fabricated on a glass substrate with circular electrodes with a radius of 2 mm, as shown in (Figure 1A and B). Both gate electrodes are circular (radius 2 mm) and are coupled to a single PEDOT:PSS channel through the same electrolyte well. The two gates address the same channel, an architecture designed to minimize device-to-device variability arising from mismatched channel impedance during fabrication and to enable direct comparison of gate-dependent responses under matched channel conditions.

**Figure 1.**
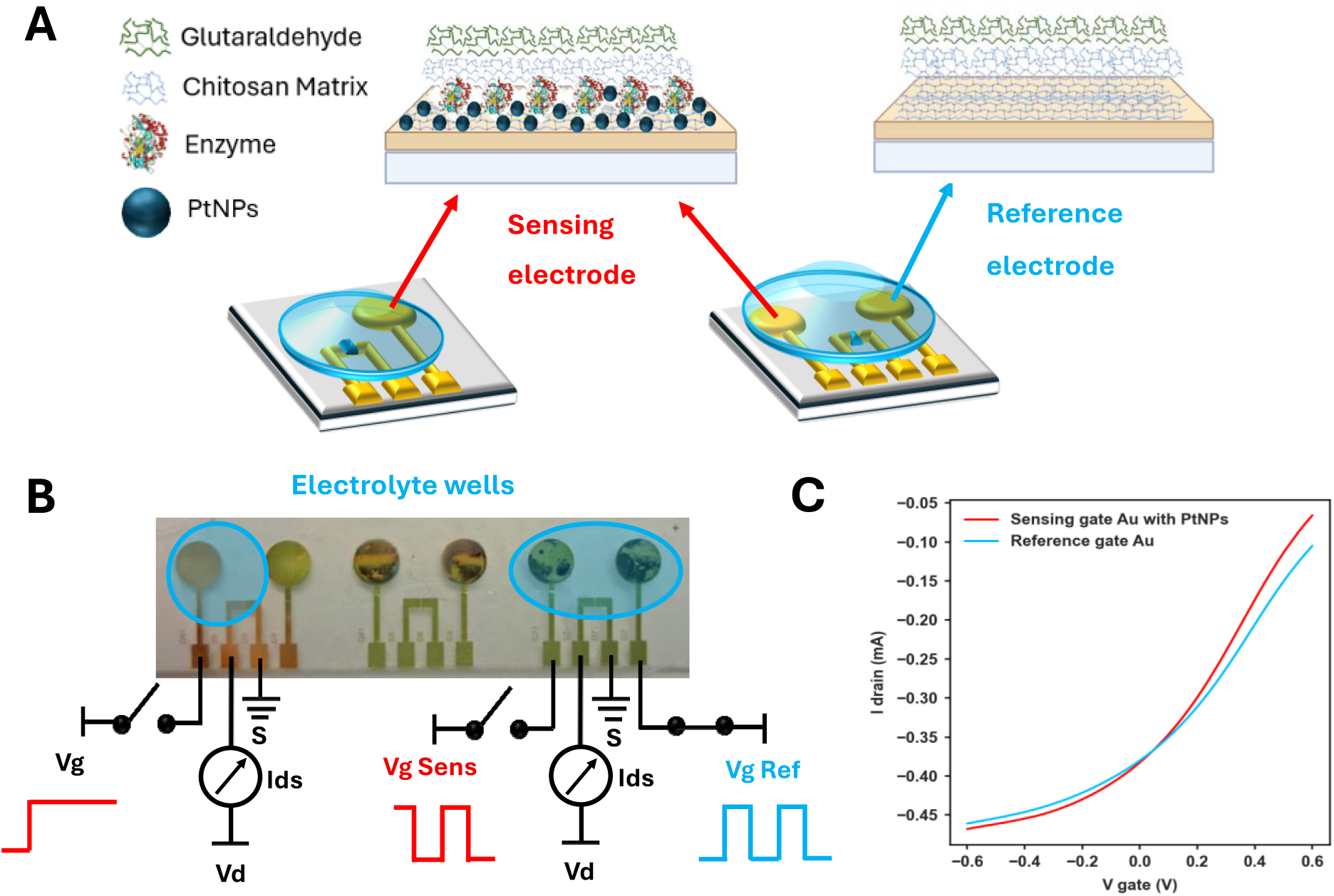
Single-gate and shared-channel dual-gate OECT architectures. **A)** Schematic comparison of the single-gate and a shared-channel dual-gate OECT configuration. The sensing gate is functionalized with Pt nanostructures and glucose oxidase (GOx) immobilized in a chitosan-glutaraldehyde matrix, while the reference gate is prepared using the same matrix but without GOx and without Pt nanostructures. **B)** Photograph of the fabricated device and corresponding schematic showing the single- and dual-gate electrolyte wells and electrical connections to the gate-voltage source. **C)** Representative transfer characteristics of the shared PEDOT:PSS channel gated independently by the sensing gate and the reference gate in PBS (pH 7.4) with V_DS_ = -0.2 V and V_G_ swept from -0.6 to +0.6 V.

The sensing gate consists of 10 nm titanium (Ti), covered by 100 nm gold (Au), modified with a platinum nanostructure layer (PtNPs), followed by the immobilization of glucose oxidase (GOx) crosslinked with a chitosan–glutaraldehyde matrix (Figure 1A). This functionalization strategy has been shown to increase the long-term stability of the enzyme in aqueous electrolytes and complex media.^26,27^

The reference gate was functionalized following the same procedure but omitting the GOx enzyme and the PtNPs to reduce the likelihood of unintended electrocatalytic reactions at the reference electrode since PtNPs alone have been shown to catalyze redox reactions (e.g., H₂O₂ oxidation) and thus can generate undesired false-positive currents and introduce gate-dependent artifacts.^28^ In the present work, the term “reference gate” therefore denotes a non-enzymatic gate electrode intended to track channel conductance state and electrolyte-level variations, positioned in the same electrolyte as the sensing gate (Figure 1B). The corresponding representative transfer curves (Figure 1C) show depletion-mode behavior typical of PEDOT:PSS OECTs: applying a positive gate bias drives cations from the electrolyte into the channel, compensating the PSS sulfonate groups, reducing the hole density in the conducting polymer, and decreasing the drain current. The corresponding transconductance traces are shown in Figure S1, Supporting information. These measurements verify that both gate electrodes can modulate the shared channel under identical channel geometry and electrolyte conditions, providing the electrical baseline for subsequent dual-gate self-calibration experiments.

### 2.2 Dual-gate system operation enables drift cancellation

To address the time-dependent baseline drift observed during continuous single-gate operation (**Figure S2a,** Supporting information), a behavior widely reported for OECTs in aqueous/physiologically relevant conditions and commonly attributed to slow evolution of the device state (e.g., ionic/redox equilibration, charge retention, and interfacial adsorption processes)^5,29,11^, we adopt an active control approach rather than relying on static biasing.

Specifically, we introduce a three-pulse protocol with a sense-replenish-ground pulsing strategy shown in **Figure 2A** that periodically reconditions the PEDOT:PSS channel state, thereby stabilizing the baseline and improving separation of analyte-dependent response from drift.^10^

**Figure 2.**
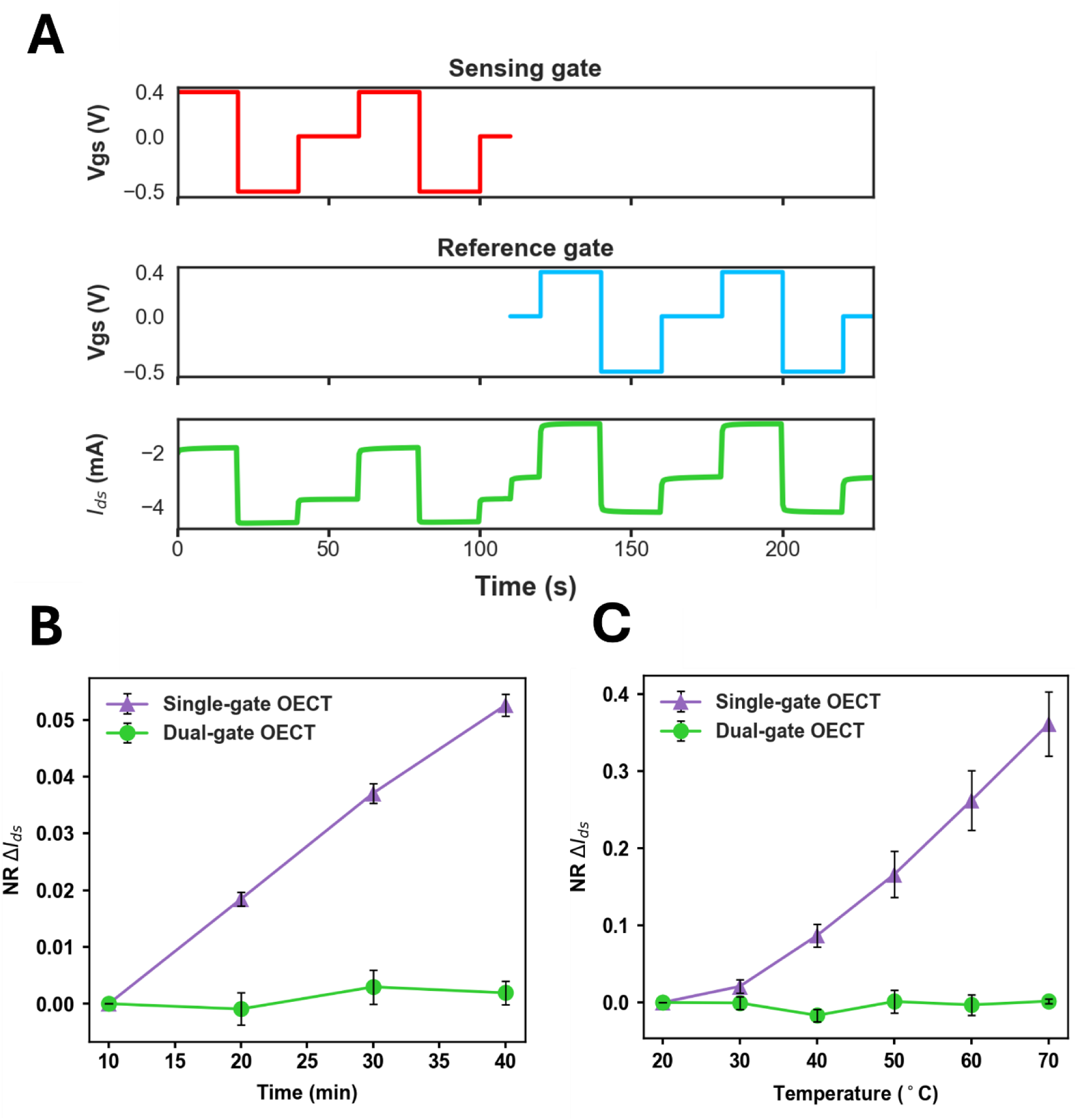
Dual-gate triple-pulse operation suppresses time- and temperature-dependent drift.A) Representative dual-gate triple-pulse protocol applied sequentially to the sensing gate and then reference gate sharing the same PEDOT:PSS channel. The sense-replenish-ground cycle V_G_ = +0.4 V`→` −0.5 V `→` 0 V is applied with a pulse width of 20 s. The analytical drain current is extracted exclusively during the sense phase at 0.4 V after a 15 s settling period by averaging the final 5 s of data. **B)** Comparison of time-dependent drift under continuous single-gate operation and after dual-gate output in the presence of 50 µM. The dual-gate OECT system (green) suppresses time-dependent drift by ∼ 96% at 40 min relative to a single-gate OECT output (purple) (Single-gate: NR = 0.053; dual-gate: NR = 0.002). Drift is calculated as 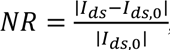, where I_ds, 0_ is defined as the pre-analyte drain current measured at 0 µM glucose. Data are mean NR ± SEM ; n = 4. **C)** Comparison of temperature-induced variation under continuous single-gate operation and under dual-gate operation during sensing in 50µM glucose. At 70 ⁰C, the single-gate deviation reached NR = 0.36, whereas the dual-gate output remains near the baseline (NR = 0.002), corresponding to approximately 99 % suppression of temperature-induced variation. Data represent NR ± SEM , n = 3 devices.

During the sense phase, the gate is stepped to *V_G_* = + 0.4 V (selected to match the optimal enzymatic glucose readout conditions used throughout this work) and the analyte-dependent drain current is recorded. The subsequent replenish step ( *V_G_* = − 0.5 *V*) is introduced to promote recovery of the channel conductivity and thereby help suppress ionic memory effects associated with prior bias history. Finally, a ground/reset step (*V_G_* = 0 *V*) is applied to relax residual ionic polarization at the gate surfaces and stabilize the baseline prior to the next cycle.

The drain current is extracted exclusively during each sense phase (only at *V_G_* = + 0.4 V), after 15 seconds settling period, by averaging the final 5 seconds of data, thereby reducing sensitivity to within-pulse transients (**Figure 2A**). The extracted currents from the repeated sense pulses are then averaged to obtain a single representative current.

Pulsing the sensing gate first allows the enzymatic reaction to reach a quasi-steady value within the extraction window before switching to the reference gate in the system. The subsequent reference-gated readout then samples the shared channel under the same post-sense operating condition, reducing contributions from within-cycle electrochemical dynamics. This strategy mitigates ion-retention artefacts and improves separation of sensing-specific modulation from common-mode channel dynamics.

Drift in OECTs can often originate from coupled evolution of channel ionic/redox processes and gate/electrolyte interfacial processes.^15–19^ In the linear operating regime, the drain current can therefore be represented using a simplified form of the Bernards-Malliaras model (Supporting information**, Equation 1**):

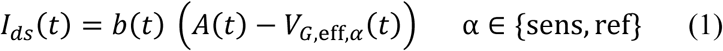

where b(t) represents a time-dependent gain term associated with the channel state and V_G,eff,**α**_(t) captures the effective gate action for the sensing or reference interface (full derivation in *Supporting information*). b(t) ∝ μC^∗^(Wd/L) depends on the charge-carrier mobility, volumetric capacitance, electrolyte properties, and bias history, which can change non-linearly with environmental conditions. Therefore, drift is inherently state-dependent and is not expected to appear as a purely additive offset with a fixed amplitude. Consequently, simple drift subtraction (e.g., I_sens_ − I_ref_) is generally insufficient.

Moreover, *Vg_eff_* is a function of analyte concentration, environmental drift and gate-surface electrochemistry. Therefore, the responses of two differently functionalised gates cannot be assumed to be perfectly common-mode, since differences in surface chemistry, such as the presence of an enzyme layer, can introduce distinct interfacial kinetics and hence different contributions to V_G,eff,_**_α_**.

Therefore, we present a reference-derived multiplicative drift-correction approach, in which the reference-gate current (*I_ref_*) is used to estimate the evolution of the shared channel drift and to dynamically rescale the sensing-gate current (*I_sens_*). Since the sensing and reference gates share the same PEDOT:PSS channel, both outputs contain a common drift component associated with channel-state evolution.

Since both gates share the same PEDOT:PSS channel, their outputs contain a common drift component associated with channel-state evolution. However, differences in gate surface chemistry and interfacial kinetics can cause this shared drift to be expressed with different amplitudes in the two readouts. We therefore describe the sensing-and reference-gate currents as coupled to the same underlying drift factor, *D* , with *α* and *β* representing its relative contribution to the sensing and reference signals, respectively **(Equation 2)**.

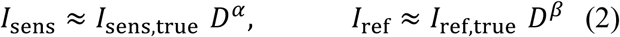

To quantify the step-to-step evolution, consecutive measurements are expressed as fractional magnitude changes, ℎ*_k_* for the sensing gate and *g_k_* for the reference gate. The reference change, *g_k_*, serves as a marker of the shared channel-state evolution, whereas ℎ*_k_* contains both the corresponding drift contribution and sensing-specific changes **(Equation 3).**

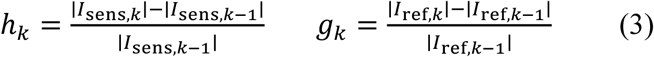

We model the sensing-gate fractional change as ℎ*_k_* ≈ ***γ*** *g_k_* + *ε_k_* , where ***γ*** is the common-mode channel coupling coefficient and *ε_k_* accounts for non-common-mode effects, including the analyte-dependent sensing response and gate-specific interfacial potential shifts.

The coefficient *γ* is a constant value, experimentally derived by least-squares fitting of the cycle-to-cycle fractional sensing-gate changes, ℎ*_k_*, against the corresponding reference-gate changes, *g_k_* (Supporting Information, Equation 18). In practice, a value of *γ* = 1 corresponds to equal drift coupling in the sensing and reference readouts, whereas *γ* > 1 indicates that *I*_sens_ is experiencing the same underlying drift stronger than *I*_ref_.

The cumulative sensing-gate correction factor *F_k_* is then updated recursively to account for the progressive accumulation of reference-tracked drift as shown in **Equation 4**.

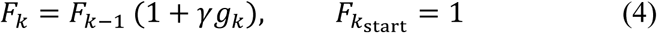

Finally, the drift-compensated dual-gate output is obtained by dividing the sensing-gate current by *F_k_* **(Equation 5).**

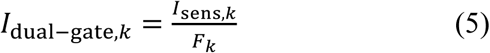

In practical terms, these expressions implement a dynamic, reference-derived gain correction: *γ* quantifies how strongly reference-tracked drift is expressed in the sensing-gate response, while *F_k_* accumulates this drift contribution over successive measurements and is used to rescale *I*_sens_. This model, combined with the *sense-replenish-ground* pulsing protocol, enables the dual-gate system to evaluate and suppress both time-dependent drift and temperature-induced variation.

Drift is reported as the magnitude-normalized response: 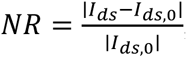, where *I_ds_*_,0_ is defined from the pre-analyte baseline in PBS (0 µM glucose). Stability was subsequently evaluated in the presence of 50 µM glucose to maintain an analyte-relevant operating background. Under continuous single-gate OECT conditions (**Figure S2a**, Supporting information), the normalized deviation increased to *NR* = 0.053 at 40 minutes, whereas the dual-gate system output remained close to the baseline (*NR* = 0.002), corresponding to a ∼96% reduction in drift magnitude at this time point (**Figure 2B**; mean NR ± SEM, *n* = 4).

The same correction strategy was then evaluated under controlled temperature variation during glucose sensing. Across 20 - 70 ⁰C, the continuous single-gate response reached *NR* = 0.36 at 70 ⁰C, while the dual-gate output remained near baseline (*NR* = 0.002), yielding a ∼99.5% suppression of temperature-induced variation during glucose sensing (**Figure 2C**; mean NR ± SEM, *n* = 3 devices). The full sensing-gate, reference-gate, and corrected dual-gate traces corresponding to the time- and temperature-variation experiments are provided in **Figure S2b),c),** Supporting information.

Together, these results indicate that combining the pulsed operation with reference-derived dynamic scaling effectively mitigates common-mode baseline evolution and stabilizes the OECT readout against both temporal drift and temperature fluctuations.

### 2.3 Dual-gate system tracks non-linear channel-state drifts, enhances recovery of the channel and improves sensitivity of analyte sensing

The OECTs are designed and functionalized to have high sensitivity in the sub-µM to low-µM glucose regime, where quantitative readout is particularly vulnerable to channel drift. **Figure S3** demonstrates that the normalized response of the reference-gated single OECT (lacking enzyme), operated in a continuous measurement mode, increases with stepwise glucose addition, indicating that the readout is influenced by channel state rather than analyte transduction. Moreover, the reference-gated OECT response increases even when glucose concentrations are added in decreasing order (from 10 mM to 10 µM; **Figure S4B**), further suggesting that the effect is not driven solely by analyte accumulation but by drift or electroactive oxidative behavior.

Control experiments further excluded chemical or electrostatic crosstalk as the origin of this behavior. Isolated Au and Pt electrodes, as well as functionalized electrodes lacking enzymatic coupling, were tested under different conditions and exhibited no glucose-dependent response (**Figure S4A ,B**), indicating that the apparent reference-gate signal arises from channel-state evolution rather than H₂O₂ diffusion, Debye length, or Pt nanoparticle-mediated redox activity through the shared electrolyte.

To further isolate the origin of the non-enzymatic OECT response, a single reference gate-only OECT was evaluated under three channel conditions: (i) reference gate in a separate electrolyte well (isolated), (ii) reference gate co-located with the sensing gate in the same well without prior channel conditioning, and (iii) the same configuration but following channel pre-conditioning by repeated bidirectional gate-voltage sweeps **(Suppl. Figures S5A and B)**. The apparent normalized response (NR) depended strongly on the channel conditioning state. The NR of 0.075, when the reference gate was isolated in a separate well after a single bidirectional gate sweep, increased to NR ≈ 0.153 when co-located in the same well without channel pre-conditioning, indicating that the magnitude of the NR is governed primarily by the channel conditioning state rather than by co-location. After a full set of bidirectional pre-conditioning sweeps, the NR decreases to ≈ 0.050 (**Figure S3A**). Thus, pre-conditioning suppresses the channel-state-associated reference-gate NR by ∼67% relative to the unconditioned co-located state, consistent with partial reversibility of channel-state evolution under bidirectional biasing.

These observations are consistent with an interpretation in which the reference-gated signal primarily reports progressive evolution of the PEDOT:PSS channel state, consistent with ionic retention/entrapment in the volumetrically doped film, rather than analyte transduction. Specifically, with repeated positive gate bias, cations may gradually accumulate in trap sites, leading to asymmetric ionic injection/extraction and retention of ionic charge within the volumetrically doped channel, reducing the effective conductivity.^30,17^ Such entrapment behavior has been documented in PEDOT:PSS and related composite films and has been deliberately exploited for designing organic electrochemical memory devices.^17,19,31,32^

In the shared-channel dual-gate configuration, the sensing- and reference-gate components show distinct concentration-dependent evolution during stepwise glucose addition (**Figure 3A**). The reference-gate response remains smaller than the single-gate reference response measured under continuous operation (**Figure S3C**, Supporting Information), but remained non-zero, indicating that residual channel-state contributions can still be monitored during dual-gate sensing. Since both gates are coupled to the same channel and the reference-gate signal provides an in-situ reporter of common-mode channel-state evolution, the reference-derived correction factor is therefore applied to rescale the sensing-gate output and generate the corrected dual-gate response shown in **Figure 3B**.

**Figure 3.**
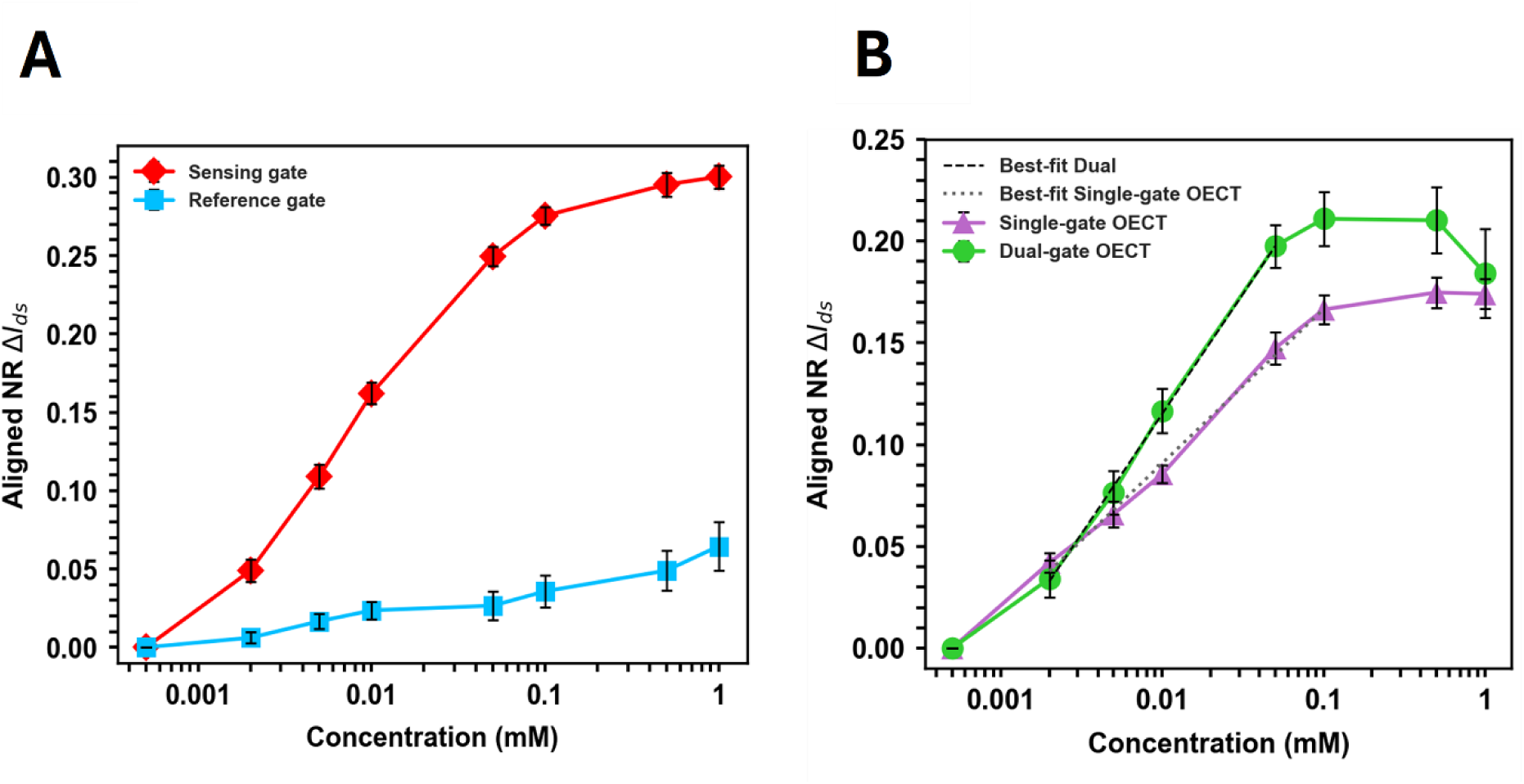
Dual-gate correction improves glucose sensitivity compared with continuous single-gate OECT operation. **A)** Dual-gate OECT signal components plotted as aligned 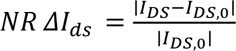 versus glucose concentration (log scale) for the sensing gate (red) and reference gate only (blue), showing channel-specific effects prominent at higher concentrations. Data are mean NR ± SEM ( n = 3 devices.) **B)** Comparison of the continuous single-gate OECT response (purple) and dual-gate output (green) versus glucose concentration (log scale). The dual-gate response is obtained from the sensing- and reference-gate signals in panel A) using the reference-derived correction factor F_k_. Dashed line indicates linear bestfit of the form ΔNR = A + S ∗ log_10_(C) over the indicated concentration range (R = 0.997 for both single-gate and dual-gate OECT). Error bars: mean NR ± SEM (n = 3 devices).

The dual-gate output is then compared directly with continuous single-gate OECT operation. The continuous single-gate OECT shows a linear response from 500 nM to 100 µM with a calibrated sensitivity of S = 7.6 % NR per decade (R = 0.997; mean NR ± SEM, n = 3 devices) (**Figure 3B, purple**). In comparison, the corrected dual-gate output shows a linear response from 500 nM to 50 µM with a steeper calibration slope of S = 11.8 % NR per decade (R = 1; mean NR ± SEM, n = 3 devices), corresponding to a ∼55 % increase in calibrated sensitivity (**Figure 3B, green**).

The enhanced sensitivity is primarily attributed to the negative replenish step, which is introduced to promote recovery of PEDOT:PSS channel conductivity after the positive sensing bias and thereby to suppress ionic memory effects associated with prior bias history. This interpretation is consistent with PEDOT:PSS-based OECT operation, where positive gate bias drives ion penetration and channel dedoping, while bias reversal promotes recovery toward the conducting state. The use of negative gate bias has previously been exploited to reduce state retention associated with asymmetric ion injection and to facilitate reversal of retained states.^³¹,¹⁷^

Consistent with these results, the dual-gate protocol improves sensitivity by actively controlling channel-state evolution, rather than simply increasing enzymatic signal amplification. This improvement occurs over a narrower optimized linear range than continuous single-gate operation, highlighting a trade-off between broad operational range and enhanced sensitivity in the targeted glucose regime.

To further validate our pulsing strategy, we incorporate the same OECT channel into a dual-gate architecture and compare two pulsing protocols: (i) bipolar gating between +0.4 V and 0 V and (ii) the full pulsing sequence of +0.4 V → -0.5 V → 0 V (Suppl. **Figures S6A and B**). Under the two-pulse protocol, the glucose sensitivity is S = 7.5% NR per decade, comparable to the value obtained for the single-gate configuration. In contrast, the triple-pulse protocol yields an increased sensitivity of S = 9.5% NR per decade over the same concentration range (Figure S5C and D; mean NR ± SEM, n = 3 devices). Importantly, the reference-gate normaliied response is significantly reduced under the triple-pulse protocol. At matched concentration conditions, the reference NR reachs ∼0.06 under bipolar pulsing, whereas it is limited to ∼0.03 under triple-pulse operation, corresponding to ∼50% suppression of the channel-state-associated artefact. These results indicate that inclusion of the negative gate enhances recovery of the PEDOT:PSS channel between gate bias steps, thereby reducing cumulative channel-state evolution. The improved sensitivity observed under triple-pulse operation is therefore attributed not to increased enzymatic activity, but to more effective control and compensation of channel-state-dependent contributions.

Most importantly, in our dual-gate system, the sensing-gate output is not treated as a concentration-only signal. Instead, the reference gate provides an in-situ proxy for common-mode channel-state evolution, where common drift is converted into a cumulative correction factor to rescale the sensing response. This allows rescaling of the sensing response and thereby more reliable isolation of the analyte-correlated component by accounting for channel- and environment-driven drift.

### 2.4 Dual-gate system restores sensitivity in the presence of interferents and complex media

The dual-gate system’s ability to decouple drift from true analyte response via the reference-gate readout is valuable for mitigating interferent-induced shifts in the OECT current output. Interfering species and complex-media-dependent ionic compositions can affect enzymatic OECT output by modifying the effective doping state of PEDOT:PSS. While ion-selective membranes, molecularly imprinted polymers, nanomaterials, and other gate barrier layers can improve selectivity at the gate interface, the PEDOT:PSS channel remains sensitive to variations in pH, ionic strength, and other environmental factors that modulate its volumetric conductivity.^¹⁵,³³,³⁴^ Consequently, acid-containing compounds (e.g., ascorbic acid, uric acid, lactic acid) can alter the local pH and change ion concentrations at the channel-electrolyte interface. This can modulate PEDOT:PSS channel conductivity and induce positive or negative offsets in **|I_ds_|** even when the gate surface is protected, rendering simple baseline subtraction unreliable when the sign of the offset is not known a priori.

In contrast, the dual-gate architecture uses the reference gate to measure these environment- and state-driven contributions and dynamically corrects the analyte sensing readout. Since the interferant and complex biological media measurements represent single-condition measurements rather than a sequentially evolving series, the reference-derived correction is implemented locally as a single step. Each test condition, *X*, is compared with an interferant-free glucose anchor, **GL** (1 mM glucose in PBS), and the resulting fractional change in the reference-gate current is defined as (**Equation 6**):

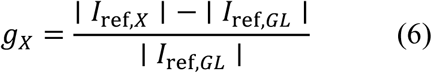

For this single-step comparison, Equations 2–5 reduce to a local correction that removes the reference-predicted common-mode contribution from the sensing-gate current magnitude (**Equation 7**; full derivations in Supporting information):

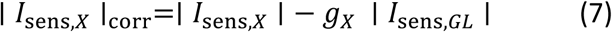

We challenged the system with high, yet physiologically relevant, interferent concentrations in PBS, i.e. 5 mM lactic acid (LA), 100 µM ascorbic acid (AA), 200 µM paracetamol (PA), and 10 mM urea (UA), to assess their effects on selectivity and PEDOT:PSS channel conductivity (**Figure 4A**). Relative to the interferent-free 1 mM glucose anchor, LA increased the raw sensing-gate **|I_ds_|**, whereas AA, PA, and UA decreased **|I_ds_|**, demonstrating that interferent-induced offsets can occur with either sign. The reference-derived calibration reduced these shifts despite high interferent levels. Quantitatively, dual-gate calibration improved **|I_ds_|** by 12–60 percentage points across interferents (**Figure S7A**; mean response ± SEM, n = 3 devices), with the largest drift reduction observed for AA (61%) and the smallest for LA (12%). This corresponds to a relative drift decrease of ∼85% for LA, ∼76% for PA, ∼96% for AA, and ∼72% for UA (**Figure S7B**).

**Figure 4.**
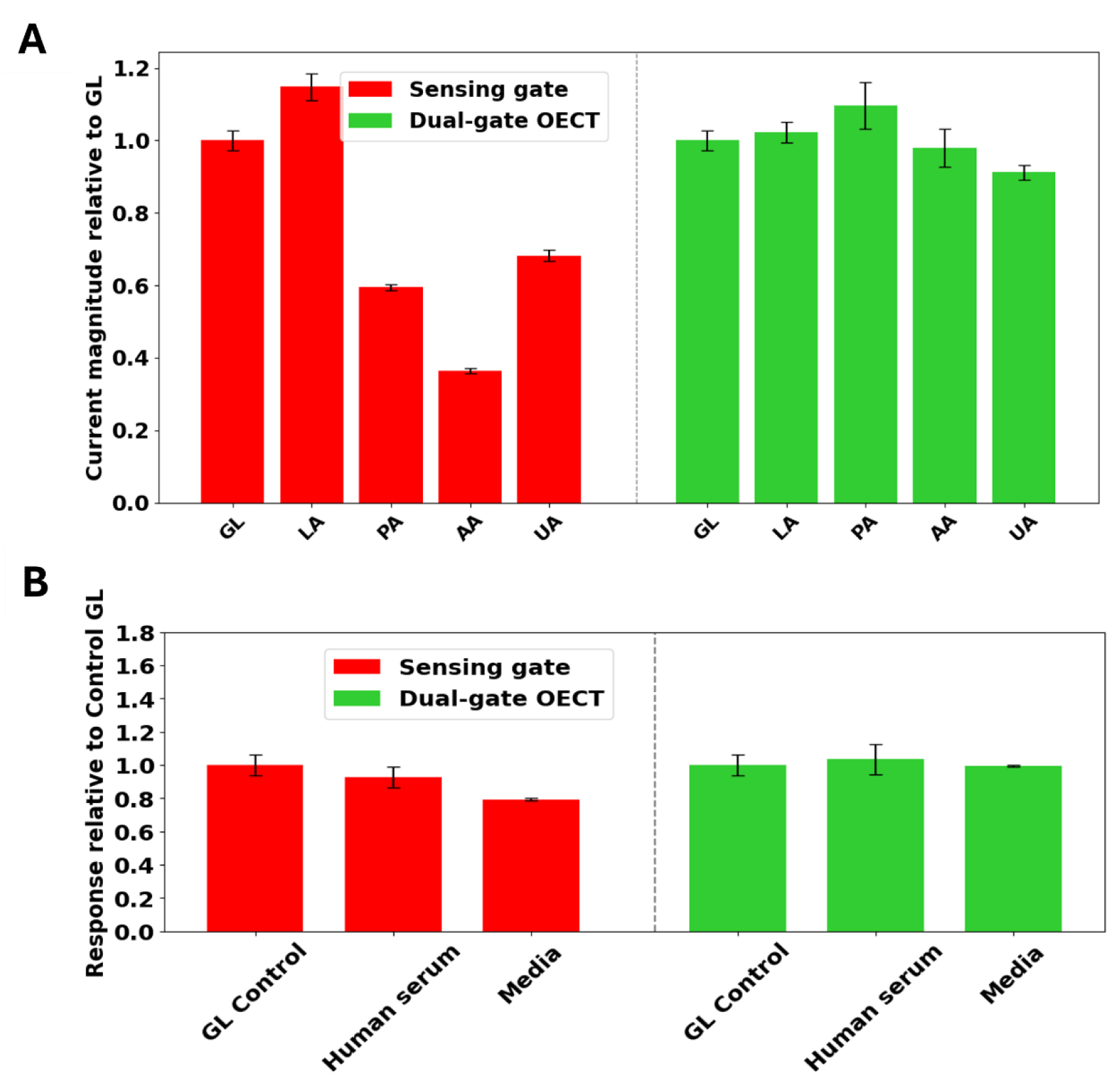
Dual-gate correction suppresses interferent- and complex-media-induced deviations during glucose sensing. **A)** Drain-current magnitude, ∣ I_ds_ ∣, measured at the +0.4 V sensing readout step for the sensing gate and reference-corrected dual-gate output at 1 mM glucose in PBS containing lactic acid, paracetamol, ascorbic acid, and urea. Concentrations are provided in the Methods. The raw sensing-gate response exhibits interferent-dependent deviations, whereas the corrected dual-gate output remains closer to the interferent-free glucose baseline. Data are mean ± SEM, n = 3 devices. **B)** Relative current magnitude for the sensing gate and corrected dual-gate output in human serum and cell-culture media, normalized to the corresponding glucose-matched PBS control. Each control contains the glucometer-measured glucose concentration of the respective complex medium and is therefore normalized to 1. The raw sensing-gate response deviates from the matched control, whereas the corrected dual-gate output shows improved agreement with the control in both matrices. Data are mean ± SEM, n = 3 devices.

Finally, we evaluate whether the dual-gate system can recover glucose readout in complex biological media. Glucose concentrations in human serum and cell-culture medium are first measured using a commercial glucometer, yielding 7.8 mM in serum and 6.0 mM in medium. For each complex medium, a PBS glucose control is prepared to match the corresponding glucometer-measured glucose concentration. For clarity, each complex-medium response is normalized to its corresponding glucometer-matched PBS glucose control. Therefore, both controls are displayed as 1 despite their different absolute glucose concentrations. When the OECT response was measured in human serum and cell-culture medium, the sensing-gate current amplitude **|I_ds_|** deviated from the matched PBS control, whereas the corrected dual-gate output showed improved agreement with the matched control in both matrices **(Figure 4B)**. Using the glucometer-matched PBS control as the reference, the dual-gate output reduced the deviation to 3.41% in serum and ≈1% in cell-culture medium (mean response ± SEM, *n* = 3 devices). Collectively, these results show that the reference-corrected dual-gate architecture compensates for interferent- and complex-media-induced channel effects, enabling drift-suppressed glucose readout in PBS, human serum, and cell-culture medium.

## 3. Conclusion

This study introduces a shared-channel dual-gate OECT architecture that mitigates PEDOT:PSS channel-state evolution and environmental variability in enzymatic sensing. By integrating an in-plane reference gate with a functionalized sensing gate addressing the same channel volume, the platform provides operando channel-state referencing while preserving biochemical transduction at the sensing interface.

Drift is addressed through a combined strategy: (i) a sense–replenish–ground pulsing cycle that actively reconditions the channel and suppresses ionic memory, and (ii) a reference-derived dynamic scaling model that compensates multiplicative baseline evolution without requiring perfectly matched interfacial kinetics. Together, these elements stabilize the OECT output against time-dependent drift and environmental perturbations, yielding strong suppression of baseline deviation under the tested conditions.

Beyond stability, the self-referenced readout improves the reliability of concentration measurements by treating the sensing-gate signal as a composite of analyte modulation and channel/environment contributions. The reference gate provides an operando proxy for common-mode channel drift that is propagated into a cumulative correction factor, enabling state-corrected rescaling of the sensing trace and more reliable isolation of the analyte-correlated component. This approach remains effective even when interferents induce offsets of either sign, underscoring the advantage of multiplicative correction over simple subtraction. Consistent performance is maintained across PBS and complex matrices, with reference-corrected outputs showing improved agreement in human serum and cell-culture medium and reduced interferent-induced deviations at physiologically representative interferent levels.

Overall, the shared-channel dual-gate strategy establishes a general framework for real-time channel-state referencing in enzymatic OECT biosensors, enabling more stable, quantitative, and reproducible readouts in complex environments and supporting translation toward point-of-care sensing. Beyond glucose detection, this platform could be extended to a wider range of enzymatic metabolites by pairing analyte-specific gate functionalization with the same channel-state correction and pulsed reconditioning principles. More broadly, the approach may provide a route to improve stability and drift tolerance in non-enzymatic OECT sensing modalities, including antibody-based assays, ion-selective interfaces, and aptamer-based biosensors, where baseline instability and interfacial drift remain central barriers to quantitative point-of-care analysis. Such improvements in drift resilience and readout accuracy are critical for translating OECT-based POCT devices from controlled laboratory settings to reliable real-time monitoring in clinical and home-care environments.

## 4. Experimental Section/Methods

### Materials

Photoresists (AZ nLOF 2035, AZ 10XT; Merck), developers (AZ 726 MIF; MicroChemicals GmbH), and parylene C dimer (SCS) were used for encapsulation; A-174 silane and Micro-90 (Fisher UK) served as adhesion and release agents. PEDOT:PSS (Clevios PH1000, Heraeus) was formulated with ethylene glycol (EG), dodecylbenzene sulfonic acid (DBSA), and (3-glycidyloxypropyl)trimethoxysilane (GOPS). Phosphate-buffered saline (PBS, pH 7.4, tablets) was used as the electrolyte. Chloroplatinic acid (H₂PtCl₆) and sulfuric acid (H₂SO₄) were used for Pt nanostructure deposition. Glucose oxidase (GOx, Aspergillus niger), lactate oxidase (LOx, Aerococcus viridans), chitosan, glutaraldehyde, and all other interferents lactic acid (LA), paracetamol (PA), ascorbic acid (AA), and urea (UA) - were obtained from Sigma-Aldrich and used as received. Human serum and cell-culture media (Merck) were tested; glucose reference measurements were obtained with commercial glucometers (SD Codefree, Sinocare). All reagents were of analytical grade and used as received unless otherwise stated.

### Device fabrication

OECTs were fabricated on glass substrates (26 × 76 mm², Knittel Glass) using a parylene peel-off process (Polyravas et al., 2019). Substrates were cleaned in acetone and isopropanol by ultrasonication and baked at 180 °C for 20 min. Gold electrodes (100 nm) with a titanium adhesion layer (10 nm) were patterned via photolithography (AZ nLOF 2035) and e-beam evaporation, followed by lift-off in acetone. A silane adhesion layer (A-174, 3%) was applied prior to deposition of two 2 µm parylene C layers separated by a Micro-90 anti-adhesive layer. Reactive ion etching defined the channels, gates, and contacts. The PEDOT:PSS channel was prepared from Clevios PH1000 mixed with ethylene glycol, dodecylbenzene sulfonic acid (DBSA), and 1 wt% GOPS, sonicated, and spin-coated in two layers at 3000 rpm with intermediate soft-bake at 80 °C. Devices were hard-baked at 120 °C for 1 h and immersed in DI water overnight to remove residual low-molecular-weight species.

### Device Functionalisation

Enzyme immobilization was performed following established procedures but was optimi**z**ed for in-vitro sensing platform [26], [27]. Platinum nanostructures were electrochemically deposited on the Au gate by chronoamperometry in 5 mM H₂PtCl₆ prepared in 50 mM H₂SO₄. A stabilizing potential of +0.7 V was applied for 1 min, followed by four cycles of −0.2 V (15 s) and +0.7 V (30 s). Subsequently, 5 µL of glucose oxidase (GOx, 4 mg mL⁻¹, Aspergillus niger) was drop-casted and dried overnight at 4 °C. A 5 mM chitosan solution was then adjusted to pH = 5.5 to promote electrostatic attachment, followed by 5 µL of 1 wt% glutaraldehyde for 15 min to crosslink the chitosan/GOx layer. Finally, the electrodes were rinsed with deionized water and dried under N₂.

### Electrical Characterisation of Device

Devices were inspected by optical microscopy to verify feature integrity and channel geometry, and insulator thickness was measured with a Dektak XT profilometer. Electrical characterisation was performed in PBS (pH 7.4) contained in a removable well, covering gate and channel. Source, drain, and gate were connected to a Keysight B1500A inside a Faraday cage; output and transfer curves were recorded under ambient conditions with a 100 ms measurement delay.

For preconditioning, bidirectional output sweeps were acquired at seven gate voltages (Vg - 0.6 - 0.6 V, in steps of 0.1 V) while sweeping Vd from -0.6 to 0 V. Bidirectional transfer characteristics were then measured by stepping Vd from -0.6 to -0.1 V with Vg swept from - 0.6 to 0.6 V (see Supplementary Figure S1). Prior to each real-time experiment, a transfer curve was collected with Vg swept from -0.6 to 0.6 V at fixed Vd = -0.1V to match the operating bias for glucose measurements.

### Continuous Operation

Continuous measurements of the drain current Ids were performed in PBS (pH 7.4) by applying a constant drain bias of 0.1V and a constant gate bias of +0.4 V, corresponding to the potential of maximum enzymatic turnover for glucose oxidation. The analyte was pipetted into the electrolyte well, and the resulting change in Ids was recorded in real time as a function of analyte concentration as shown on Suppl. Figure 3A) and B).

### Pulsing Protocol

Dynamic pulsed measurements were used to promote channel recovery and reduce bias-history-dependent drift. A three-phase sense–replenish–ground sequence was first applied to the sensing gate, with *V_G_* = +0.4V used for enzymatic readout, followed by *V_G_* = −0.5V to promote recovery of PEDOT:PSS channel conductivity and reduce ionic memory effects, and *V_G_* = 0V to relax residual interfacial polarization. The same sequence (+0.4 → −0.5 → 0 V) was subsequently applied to the reference gate, allowing the shared channel to be interrogated under the corresponding pulsed operating conditions (Figure 2A). A control two-phase sequence (+0.4 → 0 V) was used to mimic conventional enzymatic OECT operation without the replenish step (Figure S6).

### Signal extraction

At each measurement point, three repeated sense-pulse measurements were acquired. For each pulse *p*, the drain current was extracted exclusively during the +0.4V sense phase after a 15 s settling period by averaging the final 5 s:

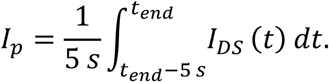

The currents were then averaged to obtain one representative current for the number of pulses

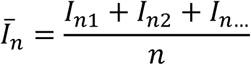

Interferents tests

Select interferents were tested at physiologically relevant upper concentrations to assess sensor specificity: 5 mM lactic acid (LA), 200 µM paracetamol (PA), 100 µM ascorbic acid (AA), and 10 mM urea (UA). Each was prepared both in 1 mM glucose PBS solution and in pure PBS to isolate individual effects. The resulting current response was compared against that of 1 mM glucose without interferents under identical pulsing conditions.

### Complex media

To evaluate sensor performance in physiologically relevant environments, glucose levels in human serum and cell culture media (Merck) were first quantified using two commercial glucometers (SD Codefree and Sinocare). The glucose concentration in human serum without added glucose was determined to be 7.8 mM (mean response, n = 3, SD Codefree), while the as-supplied culture media contained 6 mM glucose (mean response, n = 3, SD Codefree). Subsequently, PBS-based glucose solutions were prepared to match these glucometer-measured concentrations. Both serum- and media-equivalent PBS control solutions were then measured with the dual-gate system and compared with the corresponding complex media responses (Fig. 4).

## Data Availability Statement

The data that support the findings of this study are available from the corresponding author upon request.

