## Supporting Information for "A Dual-Gate OECT Platform for Drift-Suppressed and High-Sensitivity Real-Time Biosensing"

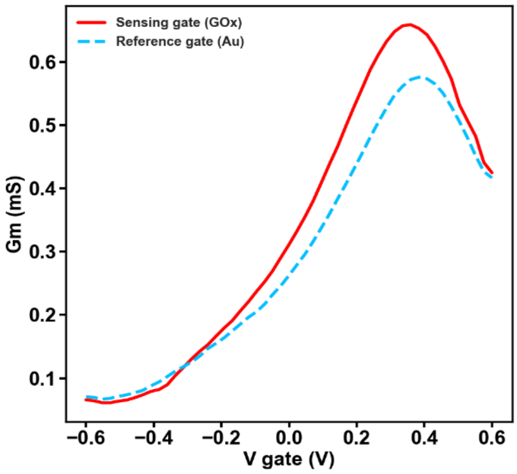


***Figure S1. Transconductance ($g_m$) characteristics** extracted from the transfer curves of the sensing and reference gates, each measured independently in a single-gate OECT configuration under the same -0.1V drain bias.*

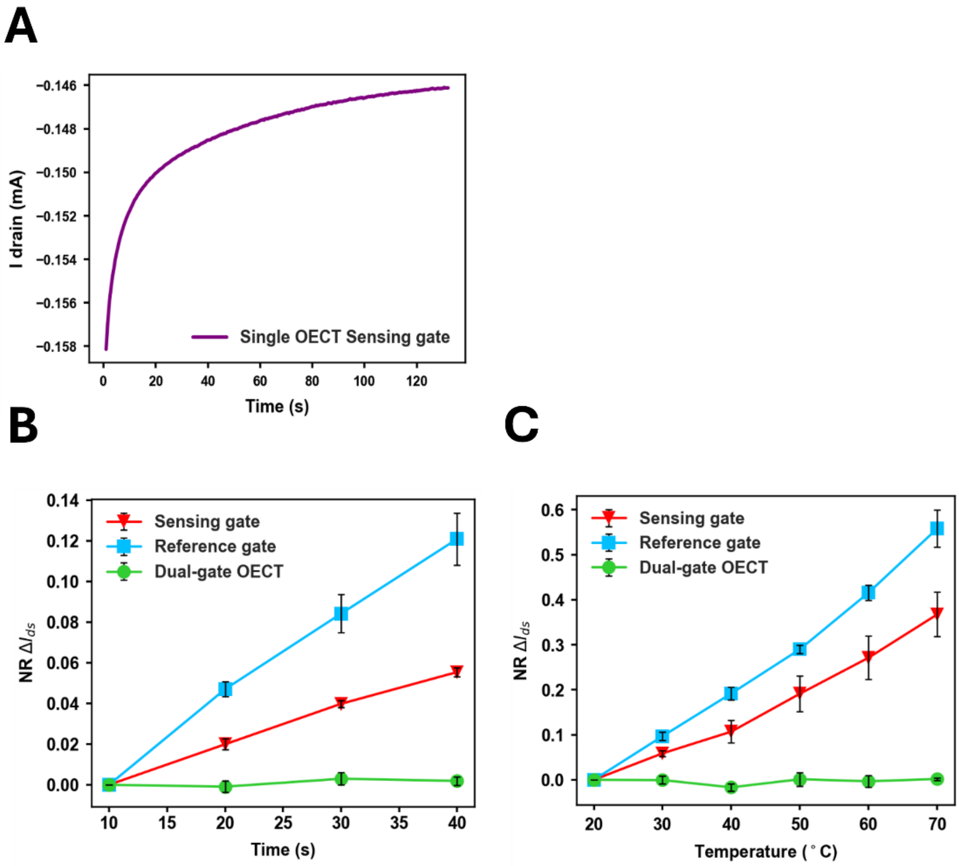


***Figure S2. Supporting traces for time- and temperature-dependent drift correction.***
***a)*** *Representative continuous single-gate OECT standard readout showing time-dependent drift of $I_{ds}$ under constant bias of $V_{DS}$ = -0.2 V and $V_G$ = 0.4 V.*
***b)*** *Dual-gate signal decomposition for the time-dependent drift experiment in 50 µM glucose, showing the extracted sensing-gate signal, reference-gate signal, and dual-gate output.*
***c)*** *Dual-gate signal decomposition for the temperature-variation experiment across 20-70 °C in 50 µM glucose, showing the extracted sensing-gate signal, reference-gate signal, and corrected dual-gate output.*
*For panels b) and c), the reported currents are extracted from the final 5 s of the sense phase following a 15 s settling period, as defined in Figure 2A.*

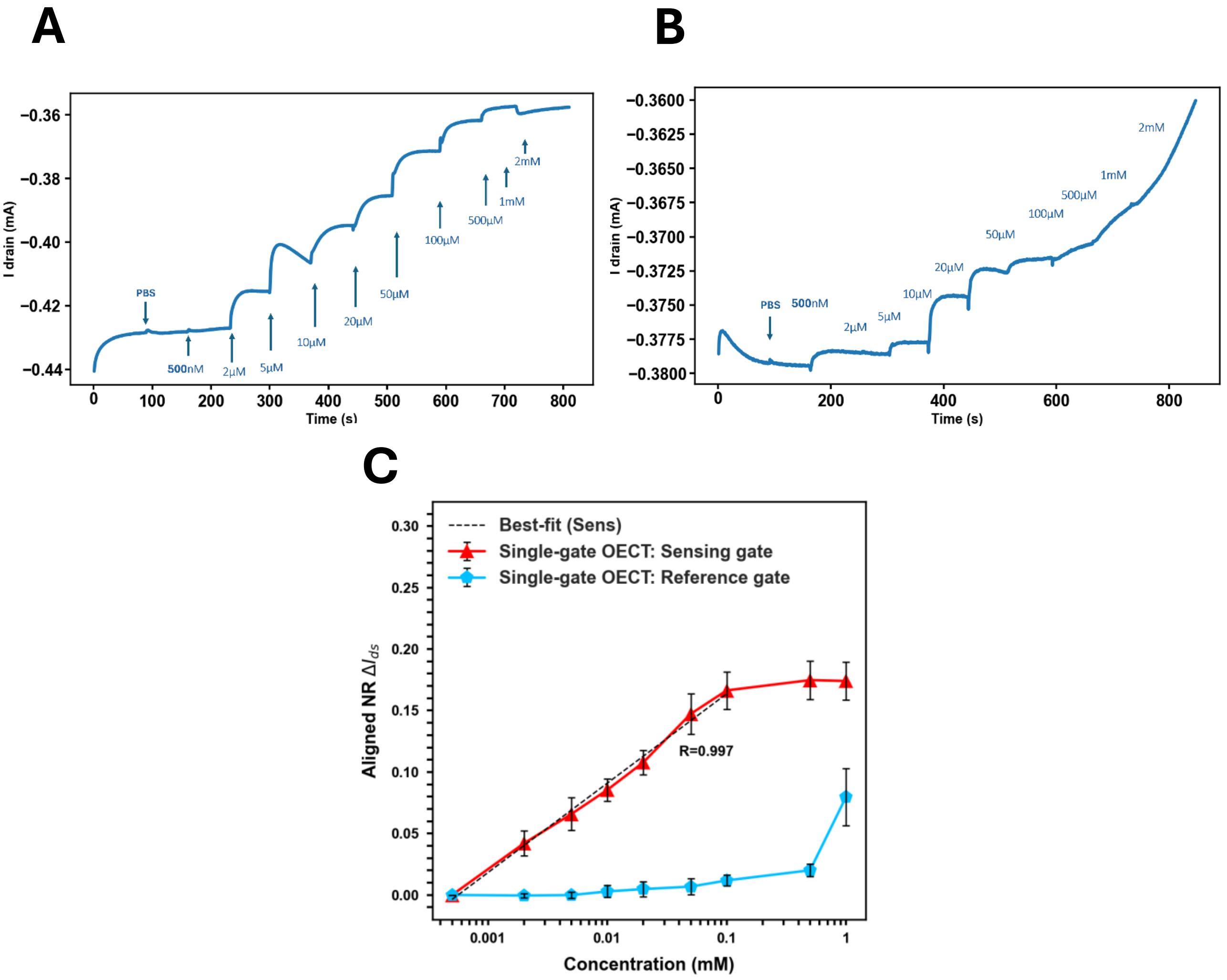


***Figure S3. Independent single-gate responses under continuous operation during stepwise glucose addition.***

*A) Representative sensing-gate response during sequential addition of glucose from 500 nM to 1 mM, showing a decrease in $I_{ds}$for the enzymatically functionalized single-gate OECT.*

*B) Representative reference-gate response under the same glucose-addition sequence, showing evolution of $I_{ds}$despite the absence of GOx.*

*C) Aligned normalized response plotted versus glucose concentration for the sensing-gate-only (red) and reference-gate-only (blue) configurations. Dashed line indicates linear best fit of the form $\Delta NR = A + S * log_{10}(C)$ over the indicated concentration range (R = 0.997).*

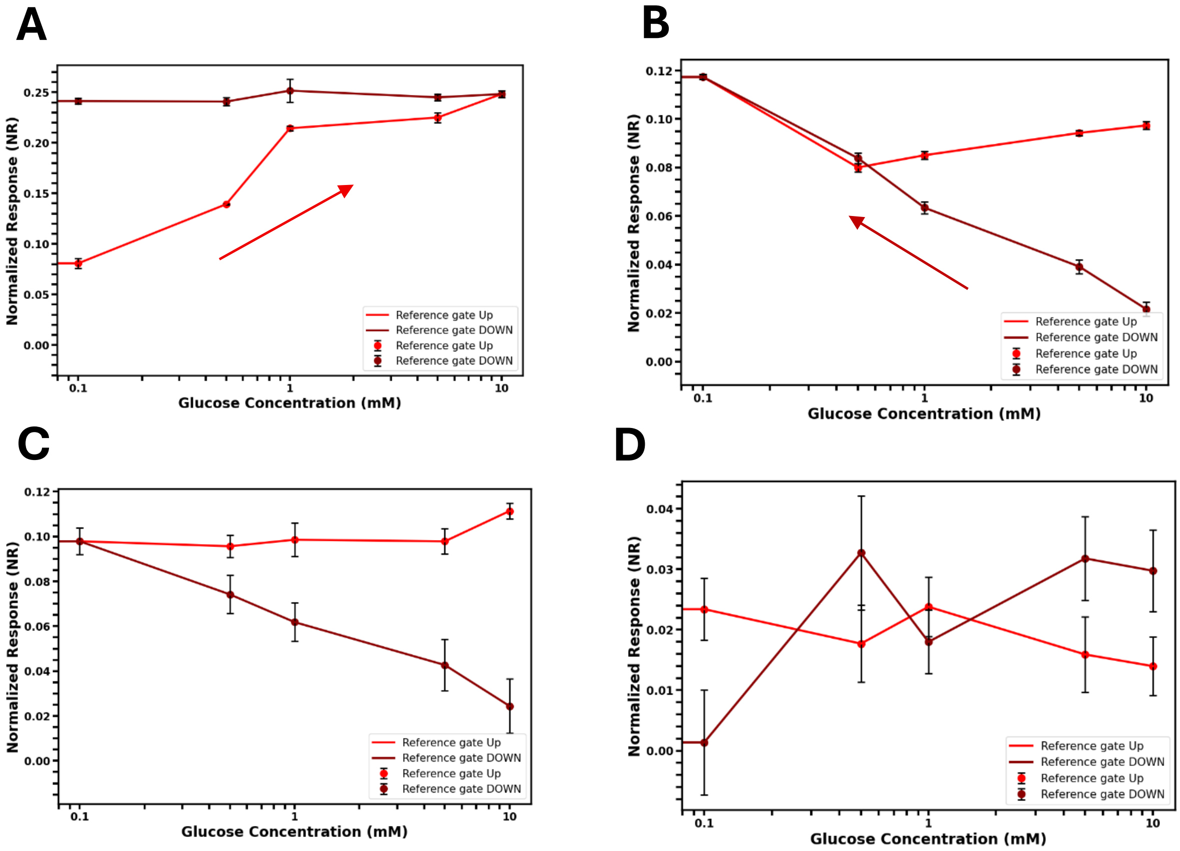


***Figure S4. Normalized response (NR) of a reference-gated OECT lacking enzyme, measured under continuous operation while the reference gate was physically separated/isolated from the sensing-gate. A)*** *Glucose is added stepwise in increasing order (red; 10 µM → 10 mM) followed by decreasing order (dark red; 10 mM → 10 µM). The arrow indicates in which direction the increasing concentrations have been added.* ***B)*** *Glucose is added stepwise in decreasing order (dark red; 10 mM → 10 µM) followed by increasing order (red; 10 µM → 10 mM). The arrow indicates in which direction the decreasing concentrations have been added.* ***C)*** *Successive additions performed without a channel pre-conditioning, using a decreasing sequence followed by an increasing sequence (dark red then red).* ***D)*** *Identical concentration sequences to panel C, but with channel pre-conditioning including a gate sweep, applied between the decreasing and increasing series. Across all conditions, the reference-gated NR increases monotonically despite reversal of the concentration order, indicating that the apparent response is dominated by operation-history/channel-state evolution rather than concentration accumulation.*

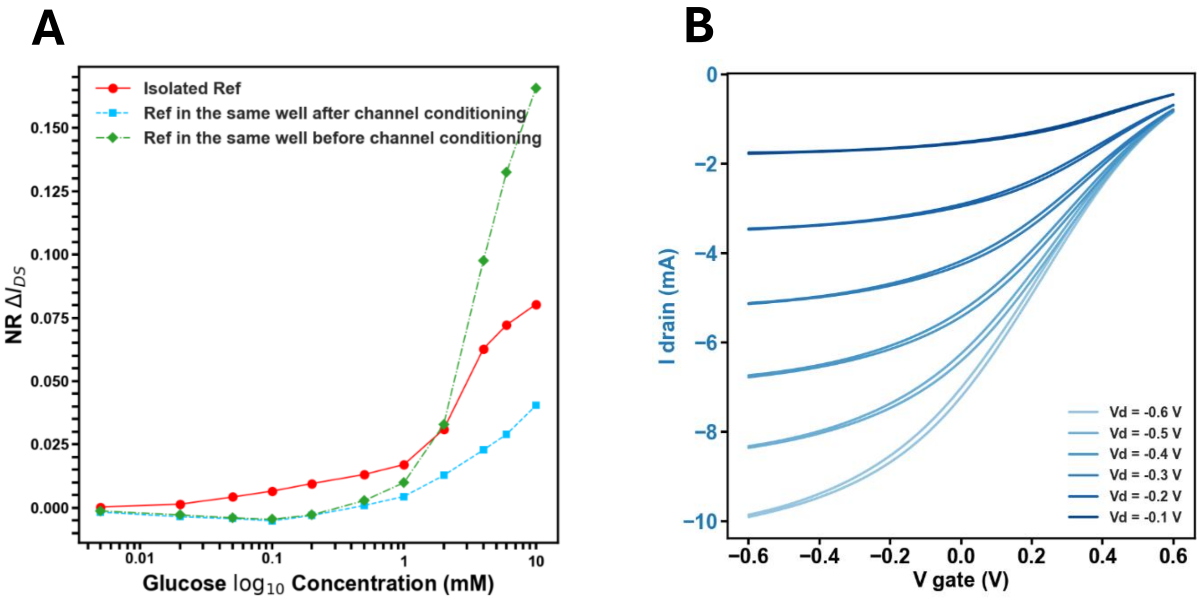


***Figure S5. Effect of channel preconditioning on the reference-gated OECT response.***
***A)*** *Single-gate OECT aligned NR output plotted versus glucose concentration (log scale) for a single-gate OECT with reference gate only, showing channel-specific effects, prominent at higher glucose concentrations. Three channel conditions are shown for the single-gate OECT: (i) reference gate is isolated from the sensing gate in a separate electrolyte well (red) preconditioned with a single sweep, (ii) reference gate is in the same well as the sensing gate but without pre-conditioning the channel (green), and (iii) in the same well as the sensing gate but pre-conditioned with a set of gate sweep cycles (blue), showing that the behavior is most likely coming from the channel state.* ***B)*** *The pre-conditioning of the channel refers to bidirectional sweeping of the gate voltage between -0.6V and 0.6V at distinct drain bias: from -0.1V to -0.6V, where single sweep refers to a gate voltage sweep under one distinct drain bias.*

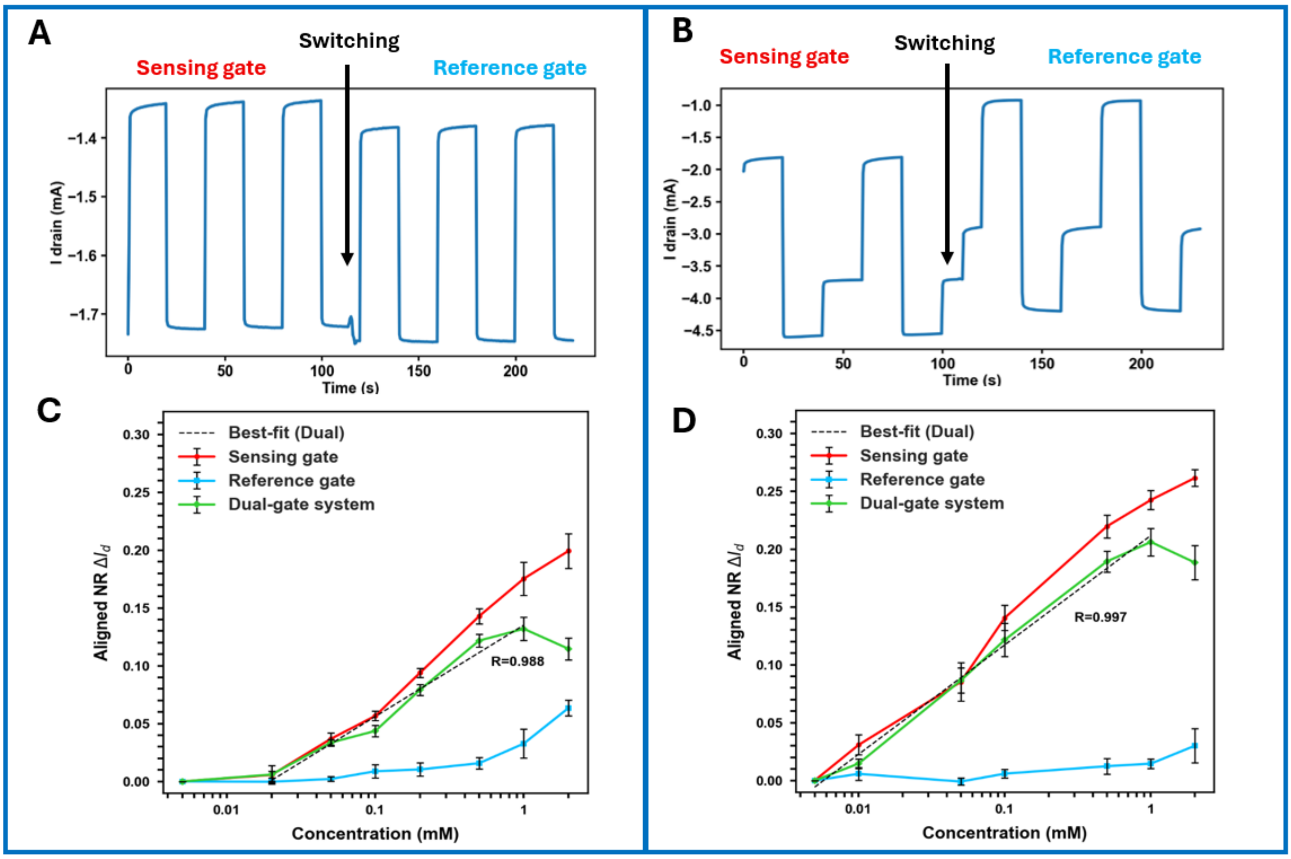


***Figure S6. Comparison of bipolar (+0.4 V ↔ 0 V) and triple-pulse (+0.4 V → –0.5 V → 0 V) operation in a shared-channel dual-gate OECT. A)*** *Protocol I (bipolar): the sensing gate is pulsed between 0 V and +0.4 V, followed by sequential pulsing of the reference gate between 0 V and +0.4 V, to mimic conventional single-gate pulsed operation.* ***B)*** *Protocol II (triple-pulse): the sensing gate is pulsed using +0.4 V → –0.5 V → 0 V, followed by the same sequence applied to the reference gate.* ***C)*** *Sensing-gate aligned NR at the +0.4 V readout step plotted versus glucose concentration (log scale) for Protocol I and Protocol II. Sensitivity S is extracted as the slope of a linear fit of NR over the indicated concentration range and reported as % NR per decade.* ***D)*** *Reference-gate NR at the +0.4 V readout step plotted versus glucose concentration (log scale) for Protocol I and Protocol II, illustrating suppression of the channel-state-associated artefact under the triple-pulse sequence. Data are mean ± SEM;* $n = 3$.

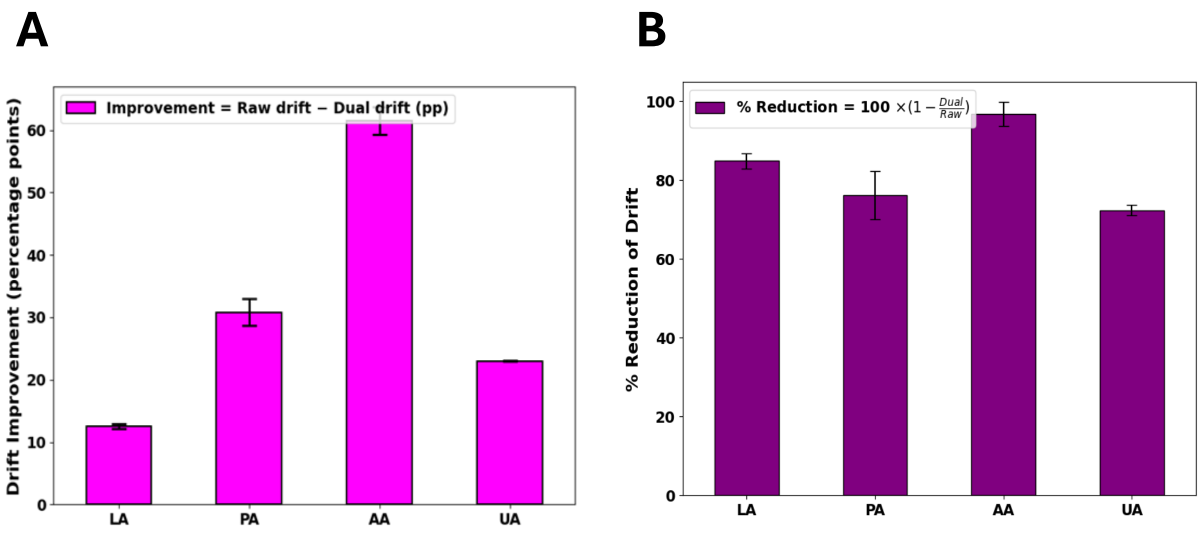


***Figure S7. Quantification of interferent-induced drift reduction after dual-gate correction.*** ***A)*** *Absolute decrease in drift, expressed in percentage points for the dual-gate system response compared to single sensing gate response. Drift is defined as the decrease in absolute percent deviation from the interferent-free glucose baseline:* $\Delta Drift(X) = Drift_{raw}(X) - Drift_{dual}(X)$, $Drift(X) = 100 \mid \frac{I_X - I_{GL}}{I_{GL}} \mid$, *where* $I_{GL}$ *is the interferent-free 1 mM glucose sensing-gate signal for the same device/replicate and* $I_X$ *is the sensing-gate signal at 1 mM glucose in the presence of the indicated interferent. Positive* ***ΔDrift*** *indicates reduced drift after correction. Values are mean % points ± SEM across devices;* $n = 3$ .

***B)*** *Relative percent reduction in drift after dual-gate correction, defined as the relative reduction in absolute deviation from baseline:* $100 \times (\mid S_{sense} - S_0 \mid - \mid S_{dual} - S_0 \mid)/\mid S_{sense} - S_0 \mid$.

**Reference-derived Multiplicative Drift Correction Derivation**

An enzymatic reaction induces a local potential drop at the surface of the electrode. This effectively shifts the gate/electrolyte potential as described in equation, resulting in an equivalent potential shift Vg eff, proportional to the amount of analyte oxidised in the electrolyte (1),(2), where the constant represents proton and oxygen variation drifts. This shift modulates the electrolyte/channel potential, dedoping the PEDOT:PSS channel and decreasing the output drain current (Ids) as described by (3). [10, 26, 27]

$$V_{G,eff,\alpha}(t) = V_{G,\alpha} + V_{off,\alpha}(t). \qquad (1)$$

$$V_{G,eff,\alpha}(t) = V_{G,\alpha} + \frac{kT(t)}{2e}(1+\delta)ln([H_2O_2]_\alpha(t)) + V_{off,\alpha}(t), \qquad (2)$$

where $k$ is Boltzmann's constant, $T$is temperature, and $e$ is the elementary charge. For the reference gate (no enzyme), $[H_2O_2]_{ref} \approx 1$ and the logarithmic term is negligible under the conditions used here.

In the Bernards–Malliaras linear regime ($| V_{DS} | \ll | V_p - V_{G,eff} |$), the drain current is:

$$I_{DS,\alpha}(t) = \frac{G(t)}{V_p(t)}\left(V_p(t) - V_{G,eff,\alpha}(t) + \frac{V_{DS}}{2}\right)V_{DS}. \qquad (3)$$

The offset term can be expanded into common-mode and gate-specific contributions:

$$V_{off,\alpha}(t) = V_0 + \Delta V_{env}(t) + \Delta V_{surf,\alpha}(t) + \Delta V_{setup}(t), \qquad (4)$$

Moreover, since Vg eff is a function of analyte concentration - $\Delta V_{chem,\alpha}(t)$, environmental/complex media drift - $\Delta V_{env}(t)$ and gate-surface electrochemistry $\Delta V_{surf,\alpha}(t)$, $\Delta V_{setup}(t)$ includes the shift due to the addition of an analyte, assuming perfectly common-mode behavior is also invalid because the reference and sensing electrodes can exhibit distinct surface potentials and reaction kinetics contributions to $V_{G,eff,\alpha}$ $(5)$.

Equivalently, the effective gate term may be written as:

$$V_{G,eff,\alpha}(t) = V_G + \Delta V_{env}(t) + \Delta V_{surf,\alpha}(t) + \Delta V_{chem,\alpha}(t), \qquad (5)$$

where $\Delta V_{chem,\alpha}(t)$denotes the analyte-driven chemical contribution.

Following Bernards–Malliaras, the prefactor $G/V_p$can be related to the OECT channel-strength factor. First:

$$\frac{G}{V_p} = \mu\, c_d\, \frac{W}{L}, \qquad (6)$$

and using $c_d \approx C^* d$ gives:

$$\frac{G}{V_p} \approx \mu C^* \frac{Wd}{L} \equiv \beta, \qquad (7)$$

where $\mu$is effective mobility, $C^*$is volumetric capacitance, and $W, L, d$ are channel width, length, and thickness. The parameter $\beta$ may evolve in time as the channel ionic/redox state evolves (drift/memory), i.e., $\beta = \beta(t)$.
Under identical channel state variables $G(t)$and $V_p(t)$(shared channel), Equation (3) yields explicit sensing- and reference-gated currents:

$$I_{sens}(t) = \frac{G(t)}{V_p(t)}\left(V_p(t) - V_{G,eff,sens}(t) + \frac{V_{DS}}{2}\right) V_{DS}, \qquad (8)$$

$$I_{ref}(t) = \frac{G(t)}{V_p(t)}\left(V_p(t) - V_{G,eff,ref}(t) + \frac{V_{DS}}{2}\right) V_{DS}. \qquad (9)$$

For subsequent drift correction, it is useful to define:

$$b(t) \equiv \beta(t)\, V_{DS}, A(t) \equiv V_T(t) + \frac{V_{DS}}{2}, \qquad (10)$$

so that the drain current can be expressed in compact form:

$$I_{DS,\alpha}(t) = b(t)\, (A(t) - V_{G,eff,\alpha}(t)). \qquad (11)$$

This emphasizes that time-dependent channel-state evolution can enter as an effective multiplicative gain term through $b(t)$, while gate/interface-dependent contributions enter through $V_{G,eff,\alpha}(t)$.

Consequently, changes in channel state can scale the measured current rather than producing a fixed additive offset, such that direct subtraction of the sensing- and reference-gate currents is not generally sufficient for drift correction.

### Reference-derived drift model and correction

The derivation below separates the measured sensing-gate response into an analyte-dependent contribution and a shared channel-state drift contribution. The reference gate is used to estimate the latter, while the analyte-dependent component is retained in the corrected sensing response.

To motivate a multiplicative correction without assuming identical sensing and reference interface kinetics, we model baseline evolution using a latent drift factor $\boldsymbol{D_k}$that scales the magnitudes of the sensing and reference readouts with potentially different strengths:

$$|I_{sens,k}| \approx |I_{sens,k}^{true}|\ D_k^{\alpha}, |I_{ref,k}| \approx |I_{ref,k}^{true}|\ D_k^{\beta}. \qquad (12)$$

Step-to-previous fractional changes (implemented in code). Drift steps are computed from magnitudes as:

$$g_k = \frac{|I_{ref,k}| - |I_{ref,k-1}|}{|I_{ref,k-1}|}, h_k = \frac{|I_{sens,k}| - |I_{sens,k-1}|}{|I_{sens,k-1}|}, \qquad (13)$$

with $\boldsymbol{g_0 = h_0 = 0}$. These are the relative step changes from one timepoint/pulse cycle to the next.

Log-step formulation and first-order approximation. In log-magnitude space, the drift steps can be written as:

$$g_k \equiv ln\,|I_{ref,k}| - ln\,|I_{ref,k-1}|, h_k \equiv ln\,|I_{sens,k}| - ln\,|I_{sens,k-1}|, \qquad (14)$$

where ≡denotes "defined as." For modest step sizes, the log-step satisfies the first-order approximation $\boldsymbol{\Delta ln\,Y \approx \Delta Y/Y}$, which motivates using the fractional-step definitions in Equation (13) as the practical implementation. This is why both the log-step and linear-step methods can yield similar results when step-to-step changes are not large.

Why $\boldsymbol{h_k \approx \gamma g_k}$. If the "true" terms $|I_{sens}^{true}|$and $|I_{ref}^{true}|$are approximately constant between adjacent points compared with drift, then:

$$\Delta ln\,|I_{sens}| \approx \alpha\,\Delta ln\,D, \Delta ln\,|I_{ref}| \approx \beta\,\Delta ln\,D, \qquad (15)$$

and eliminating $\boldsymbol{\Delta ln\,D}$yields $\boldsymbol{\Delta ln\,|I_{sens}| \approx (\alpha/\beta)\Delta ln\,|I_{ref}|}$. Under the first-order correspondence between log-steps and fractional steps, this motivates:

$$h_k \approx \gamma\, g_k + \varepsilon_k, \gamma \approx \alpha/\beta, \qquad (16)$$

where $\varepsilon_k$ captures non-common-mode contributions the analyte-dependent sensing response and gate-specific interfacial potential shifts.

Log-space separation into shared drift and gate-specific terms

Using the factorized representation $I_{D,\alpha}(t) = b(t)\, K_\alpha(t)$with $K_\alpha(t) \equiv A(t) - V_{G,eff,\alpha}(t)$, the log-step can be decomposed as:

$$\Delta ln \mid I_\alpha \mid = \Delta ln \mid b \mid + \Delta ln \mid K_\alpha \mid, \qquad (17)$$

so that $g_k = \Delta ln \mid b \mid + \Delta ln \mid K_{ref} \mid$and $h_k = \Delta ln \mid b \mid + \Delta ln \mid K_{sens} \mid$, and differences in operating point/interface sensitivity can be absorbed into the proportionality factor $\gamma$and residual $\varepsilon_k$in Equation (16).

Estimation of the drift-coupling coefficient $\gamma$ and calculation of the correction factor $F_k$

The currents obtained from the repeated sense pulses are averaged to yield one representative sensing-gate current, $I_{sens,k}$, and one representative reference-gate current, $I_{ref,k}$, for measurement $k$. Thus, $k$ denotes the sequential measurement step used in the drift analysis; for a concentration series, this corresponds to the concentration step..

Fractional changes in current magnitude between consecutive representative measurements were calculated according to Equation 13, yielding $h_k$ for the sensing-gate output and $g_k$ for the reference-gate output.

The coupling coefficient $\gamma$ was estimated by least-squares fitting of $h_k$ against $g_k$ through the origin over the index set $K$:

$$\gamma = \frac{\sum_{k\in K} g_k\, h_k}{\sum_{k\in K} g_k^2} \qquad (18)$$

where $K$ denotes $k| \; t_{k-1} > 0 \; and \; t_{k-1} > 0$, the set of consecutive representative measurement transitions included in the fit. The summations in Equation 18 combine the paired $g_k$ and $h_k$ values used in the regression and do not represent a cumulative integration over time. The through-origin fit reflects the assumption that, in the absence of a fractional change in the reference-gate signal, no reference-derived common-mode correction is applied. Once estimated, $\gamma$is held fixed throughout the correction.

The cumulative correction factor $\boldsymbol{F_k}$ is initialized at the start of the correction window and propagated recursively as:

$$\boldsymbol{F_k = F_{k-1}(1 + \gamma g_k)}, \qquad \boldsymbol{F_{k_{start}} = 1} \qquad (19)$$

where $\boldsymbol{k_{start}}$ denotes the first first measurement step used for drift correction.
The roles of $\boldsymbol{\gamma}$ and $\boldsymbol{F_k}$ are distinct: $\boldsymbol{\gamma}$ is a fixed coupling coefficient describing how strongly shared drift is expressed in the sensing-gate response relative to the reference-gate response, whereas $\boldsymbol{F_k}$ is the step-dependent factor that accumulates the reference-predicted drift over successive measurements up to measurement $\boldsymbol{k}$. $\boldsymbol{F_k}$ is initialized to 1 at the beginning of the correction window and changes with $k$ because each new reference step, $g_k$, contributes an additional fractional drift increment.

Finally, the drift-compensated dual-gate output was calculated by scaling the sensing-gate current by the cumulative correction factor:

$$\boldsymbol{I_{dual,k} = \frac{I_{sens,k}}{F_k}} \qquad (20)$$

This formulation treats the reference gate as a real-time reporter of common-mode channel-state evolution while allowing the strength of this drift contribution to differ between the sensing and reference readouts.

Drift magnitude is reported as the magnitude-normalized deviation:

$$\boldsymbol{NR(k) = \frac{|\, I_k - I_0 \,|}{|\, I_0 \,|}} \qquad (21)$$

Interferants and Complex Media Correction: Single-step Implementation of the Reference-derived Multiplicative Drift Correction

To apply the step-based multiplicative drift framework (Equations 12–13) to **individual-condition experiments** (interferants and complex media), a glucose operating-point anchor is defined, “GL”, corresponding to a glucose-only condition without interferants. Each test condition $X$ is then represented as a single step relative to this anchor rather than as part of a sequential multi-step series.

Consistent with Equation (13), the local reference-gate change between GL and condition $X$ is defined as:

$$\boldsymbol{g_X} = \frac{|\boldsymbol{I_{ref,X}}| - |\boldsymbol{I_{ref,GL}}|}{|\boldsymbol{I_{ref,GL}}|} \qquad (22)$$

Under the latent multiplicative drift model (Equation 12), the same local drift factor implies

$$|\boldsymbol{I_{ref,X}}| \approx |\boldsymbol{I_{ref,X}^{true}}|\ \boldsymbol{D_X^{\beta}},\ |\boldsymbol{I_{sens,X}}| \approx |\boldsymbol{I_{sens,X}^{true}}|\ \boldsymbol{D_X^{\alpha}} \qquad (23)$$

Since only one **GL→ $\boldsymbol{X}$** transition is available for each condition, a separate condition-specific coupling coefficient is not fitted. Instead, the measured fractional reference change is mapped locally onto the sensing-gate magnitude current at the GL operating point to estimate the corresponding common-mode change in sensing-current magnitude:

$$\boldsymbol{\Delta}\ |\boldsymbol{I_{sens}}|_{\boldsymbol{cm}}\ (\boldsymbol{X}) \approx \boldsymbol{g_X}\ |\boldsymbol{I_{sens,GL}}| \qquad (24)$$

This corresponds to a first-order (small-step) approximation of the multiplicative drift effect around the anchor, and avoids fitting a condition-specific coupling coefficient from a single GL→ $X$ transition.

The reference-predicted common-mode contribution is then removed from the measured sensing-gate current:

$$|\boldsymbol{I_{sens,X}}|_{\boldsymbol{corr}} = |\boldsymbol{I_{sens,X}}| - \boldsymbol{g_X}\ |\boldsymbol{I_{sens,GL}}| \qquad (25)$$

The correction is applied independently to each replicate using its corresponding GL sensing- and reference-gate values before averaging across replicates.

Drift and improvement metrics

To quantify interference/complex media effects relative to the glucose operating point, we compute the absolute percent deviation from the anchor:

$$\boldsymbol{d_{sens}(X)} = \left| \frac{|\boldsymbol{I_{sens,X}}| - |\boldsymbol{I_{sens,GL}}|}{|\boldsymbol{I_{sens,GL}}|} \right| * \mathbf{100}, \boldsymbol{d_{dual}(X)} = \left| \frac{|\boldsymbol{I_{sens,X}}|_{corr} - |\boldsymbol{I_{sens,GL}}|}{|\boldsymbol{I_{sens,GL}}|} \right| * \mathbf{100} \qquad (26)$$

Improvement can be reported either as a difference in drift (percentage points):

$$\boldsymbol{\Delta d(X)} = \boldsymbol{d_{sens}(X)} - \boldsymbol{d_{dual}(X)} \qquad (27)$$

or as percent drift reduction:

$$\%\,\boldsymbol{reduction(X)} = \mathbf{100}\left(\mathbf{1} - \frac{\boldsymbol{d_{dual}(X)}}{\boldsymbol{d_{sens}(X)}}\right) \qquad (28)$$

Equations (22-26) implement a reference-derived, operating-point linearization of the multiplicative drift model (Equation 12): the reference gate provides the local fractional drift step $\boldsymbol{g_X}$, which is converted into a predicted common-mode shift of the sensing magnitude and subtracted to recover a glucose-referenced sensing response without imposing $\boldsymbol{\alpha} = \boldsymbol{\beta}$.
